# Genome-wide, integrative analysis implicates circular RNA dysregulation in autism and the corresponding circular RNA-microRNA-mRNA regulatory axes

**DOI:** 10.1101/712331

**Authors:** Yen-Ju Chen, Chia-Ying Chen, Te-Lun Mai, Chih-Fan Chuang, Sachin Kumar Gupta, Laising Yen, Yi-Da Wang, Trees-Juen Chuang

## Abstract

Circular RNAs (circRNAs), a class of long non-coding RNAs, are known to be enriched in mammalian brain and neural tissues. While the effects of regulatory genetic variants on gene expression in autism spectrum disorder (ASD) have been widely reported, the role of circRNAs in ASD remains largely unknown. Here, we performed genome-wide circRNA expression profiling in post-mortem brains from individuals with ASD and controls and identified 60 circRNAs and three co-regulated modules that were perturbed in ASD. By integrating circRNA, microRNA, and mRNA dysregulation data derived from the same cortex samples, we identified 8,170 ASD-associated circRNA-microRNA-mRNA interactions. Putative targets of the axes were enriched for ASD risk genes and genes encoding inhibitory postsynaptic density (PSD) proteins, but not for genes implicated in monogenetic forms of other brain disorders or genes encoding excitatory PSD proteins. This result reflects the previous observation that ASD-derived organoids exhibit overproduction of inhibitory neurons. We further confirmed that some ASD risk genes (*NLGN1*, *STAG1*, *HSD11B1*, *VIP*, and *UBA6*) were indeed regulated by an upregulated circRNA (circARID1A) via sponging a downregulated microRNA (miR-204-3p) in human neuronal cells. We provided a systems-level view of landscape of circRNA regulatory networks in ASD cortex samples. We also provided multiple lines of evidence for the functional role of ASD for circRNA dysregulation and a rich set of ASD-associated circRNA candidates and the corresponding circRNA-miRNA-mRNA axes, particularly those involving ASD risk genes. Our findings thus support a role for circRNA dysregulation and the corresponding circRNA-microRNA-mRNA axes in ASD pathophysiology.

---

Autism spectrum disorder (ASD) [Mendelian Inheritance in Man (MIM) 209850] is a heritable, complex, highly pervasive neurodevelopmental disorder that is characterized by limited social communication, restricted and ritualized interests, and repetitive behavior [1, 2]. The characteristics of ASD-related individuals affect their family and friends, and consequently burden their expenditures, schools, and society. Hundreds of genes affected by a variety of genomic variants have been reported to be associated with the etiology of ASD [3–5]. These discoveries have provided valuable biological insights into the disorder. However, the contribution of these genetic factors to this complex disease is highly heterogeneous [6], with each factor accounting for a very low percentage of the general ASD population. In addition to genetic factors, environment and gene–environment interactions have been widely demonstrated to be associated with the development of ASD [7, 8], but the underlying molecular mechanisms remain poorly understood [9]. Recently, using samples from post-mortem ASD and control brains, several studies have reported a wide range of dysregulation of gene expression [10–12] in ASD. Moreover, epigenetic factors such as alternatively spliced transcripts [11–13], long non-coding RNAs [12, 14], microRNAs (miRNAs) [15, 16], DNA methylation [17], histone trimethylation [18] and acetylation [19] were also implicated in ASD. Although these analyses based on transcriptome and genome sequencing data have provided important insights into the effects of regulatory genetic variants [11, 20], the role of co-/post-transcriptional mechanisms, which are critical to the understanding of dysregulation of gene expression in ASD, remains largely unknown. Illuminating the underlying molecular mechanisms in the etiology of ASD awaits further investigation.

Circular RNAs (circRNAs) are a class of long non-coding RNAs produced by pre-mRNA back-splicing, which generates a structure of covalently closed loop [21] of single-strand, non-polyadenylated circular molecules [22–24]. In general, circRNAs are more stable than their corresponding co-linear mRNA isoforms [23–26]. They are often expressed in temporal- and spatial-specific manners [22, 24, 27] and are especially abundant in neural tissues [28, 29]. The best understood function of circRNAs is the regulatory role of miRNA sponges [23, 30], suggesting a previously underappreciated regulatory pathway in the form of circRNA-miRNA-mRNA axes. Several circRNAs are known to play important roles in development of nervous system [29, 31, 32]. For example, CDR1as/ciRS-7, the best-studied circRNA, is highly expressed and serves as a sponge for miRNA-7 in neuronal tissues [30], and is implicated in the pathogenesis and progression of neurological diseases [33, 34]. A recent study also reported that the interactions of circEFCAB2 with miR-485-5p and circDROSHA with miR-1252-5p were highly correlated with the expression of epilepsy-associated genes CLCN6 and ATP1A2, respectively [35]. These studies clearly indicate an important role of circRNA in epigenetic control over gene expression in neural tissues. While the large-scale transcriptome analysis of miRNA-mRNA regulatory interactions have been reported [15, 36] in ASD, the role of circRNA-miRNA-mRNA regulatory axes in ASD has never been reported and remains largely unknown.

In this study, we investigated the potential regulatory role of circRNAs in ASD. Since circRNAs lack polyA tails, RNA-seq data derived from polyA-selected RNAs naturally are deprived of circRNA population. This greatly hampers the large-scale study of circRNAs in ASD. The recently released Synapse database [12], which provided a large brain sample size of both small RNA-seq data and RNA-seq data from total RNAs (rRNA-depleted RNAs without polyA-selection) of ASD cases and controls, brought an unprecedented opportunity for us to genome-wide investigate circRNA dysregulation in ASD and the corresponding ASD-associated circRNA-miRNA-mRNA regulatory axes. We performed genome-wide circRNA expression profiling in post-mortem brains from ASD patients and non-ASD controls and thereby identified 60 circRNAs and three co-regulated modules that were perturbed in ASD. We showed a shared pattern of circRNA dysregulation in the majority of ASD samples. Furthermore, by combining the miRNA dysregulation data previously detected in the same ASD and control cohort used in the current study [15], we constructed 8,170 ASD-associated circRNA-miRNA-mRNA interactions according to the common target miRNAs of the circRNAs and mRNAs. The targets of the interactions were particularly enriched for ASD risk genes, but not for genes implicated in monogenetic forms of other brain disorders. Furthermore, we selected a circRNA-miRNA axis (circARID1A-miR-204-3p axis) that was predicted to be an upstream regulator of several ASD risk genes and indeed confirmed the corresponding circRNA-miRNA-mRNA regulatory relationships in normal human astrocytes (NHAs) or primary human neural progenitor cells (hNPCs). Our findings thus help to unveil the molecular mechanism of aberrant circRNA expression and the corresponding circRNA-miRNA-mRNA regulations in idiopathic ASD.

## Results

### Identification of differentially expressed circRNAs in ASD brain

We retrieved the high-throughput RNA sequencing (RNA-seq) data of 236 post-mortem samples of frontal cortex (FC) (Brodmann area 9), temporal cortex (TC) (Brodmann area 22, 41, and 42), and cerebellar vermis (CV) from 48 individuals with ASD and 49 non-ASD controls (the Synapse database; Table 1) [12]. The RNA-seq reads examined here were derived from total RNAs with rRNA depletion, not polyA-selected RNAs, enabled us to detect circRNAs in these ASD and control samples. After removing samples with low read depth, we then used NCLscan, which was reported to exhibit the greatest precision among currently available circRNA-detection tools [37–40], to identify circRNAs and identify differential expressed species in ASD vs. control samples. We first found no significant difference in the number of identified circRNAs per million mapped RNA-seq reads between ASD and non-ASD control samples, regardless of the brain region (Fig. 1a and Additional file 1: Table S1). To increase the stringency of sample consistency, a sample was not considered in the following analysis if number of the identified circRNAs of this sample was one standard deviation below the mean of the sample set. After that, 202 samples (73 FC, 61 TC, and 68 CV samples) were retained. There were 53,427 NCLscan-identified circRNAs in the 202 samples (Additional file 2: Table S2). We evaluated the expression levels of circRNAs using the number of supporting circRNA junction reads per million uniquely mapped reads (RPM). Since several independent transcriptomics studies of ASD indicated that the pathophysiology was present in cortex [41, 42] and that the changes in transcriptomic profiles were stronger in the cortex than in the cerebellum [12], our following analysis focused on the 134 samples of the prefrontal and temporal cortex (73 samples from 45 ASD cases and 61 samples from 43 controls; Fig. 1b). Of the 36,624 circRNAs identified in the 134 cortex samples, 40% (14,775 out of 36,624) were observed in one sample only (Additional file 2: Table S2). To minimize potentially spurious events, we only considered the circRNAs observed in more than 50% of the 134 samples (Fig. 1c). A total of 1,060 circRNAs were retained (Additional file 3: Table S3), of which 99.6% (1,056 out of 1,060) have been previously identified and collected in well-known circRNA databases (circBase [43] or CIRCpedia [44]) and 99.1% (1,050 out of 1,060) were observed in human brain (Fig. 1c). Of note, 61.4% (651 events) and 47% (498 events) of the 1,060 circRNAs were also detected in mouse and mouse brain, respectively (Fig. 1c and Additional file 3: Table S3), indicating the high conservation rate of circRNA expression between human and mouse. We performed principal component analysis (PCA) and showed that circRNA expression profiles of these 1,060 circRNAs were very similar between two types of cortex regions but were quite distinct from that of cerebellum (Fig. 1d). This result is consistent with the patterns previously observed for mRNAs [11, 45] and miRNAs [15] between these brain regions. We then performed a linear mixed effects (LME) model to detect differentially expressed (DE)-circRNAs between ASD and non-ASD samples with controlling for potential confounding factors such as sex, age, brain region (FC or TC), RNA quality (RNA integrity number; RIN) host gene expression, and so on (see Methods). We thereby identified 60 DE-circRNAs (*P* < 0.05 and |log_2_(fold change)| ≥ 0.5), in which 22 were upregulated and 38 were downregulated in ASD cortex (Figs. 1b, 1e, and Additional file 3: Table S3). We found that the fold changes for the 60 DE-circRNAs were concordant between the FC and TC (Pearson’s correlation coefficient *R* = 0.75, *P* < 10^−11^; Fig. 1f).

**Table 1.**
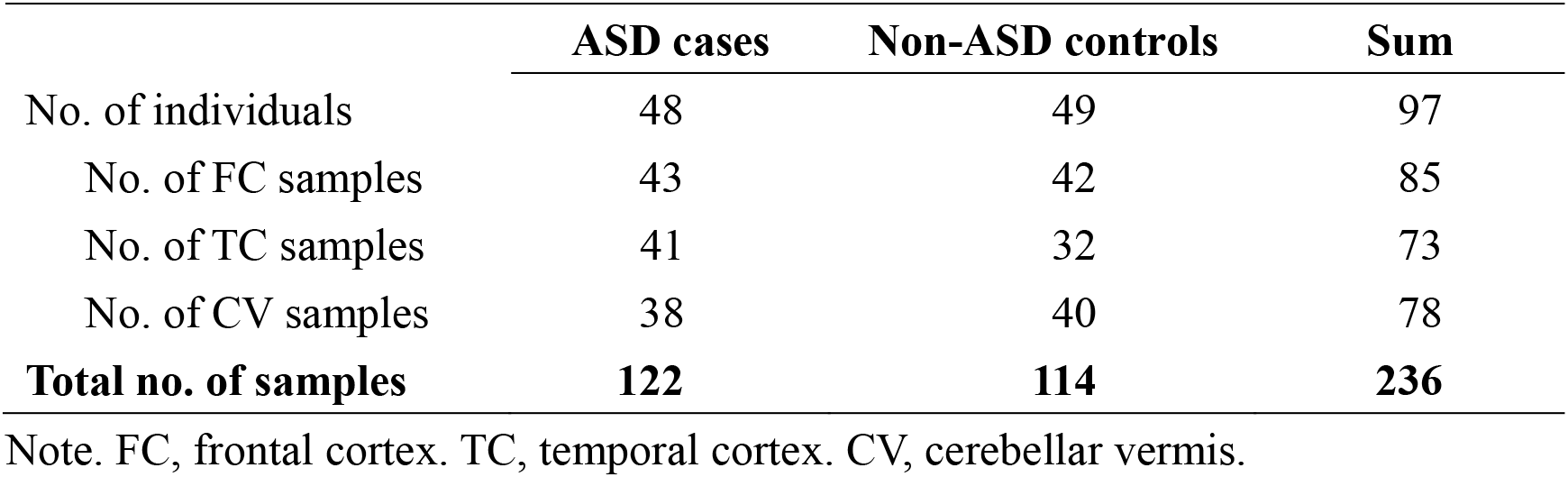
Brain tissue samples used in this studies.

|  | **ASD cases** | **Non-ASD controls** | **Sum** |
| --- | --- | --- | --- |
| No. of individuals | 48 | 49 | 97 |
| No. of FC samples | 43 | 42 | 85 |
| No. of TC samples | 41 | 32 | 73 |
| No. of CV samples | 38 | 40 | 78 |
| **Total no. of samples** | **122** | **114** | **236** |

Note. FC, frontal cortex. TC, temporal cortex. CV, cerebellar vermis.

**Figure 1.**
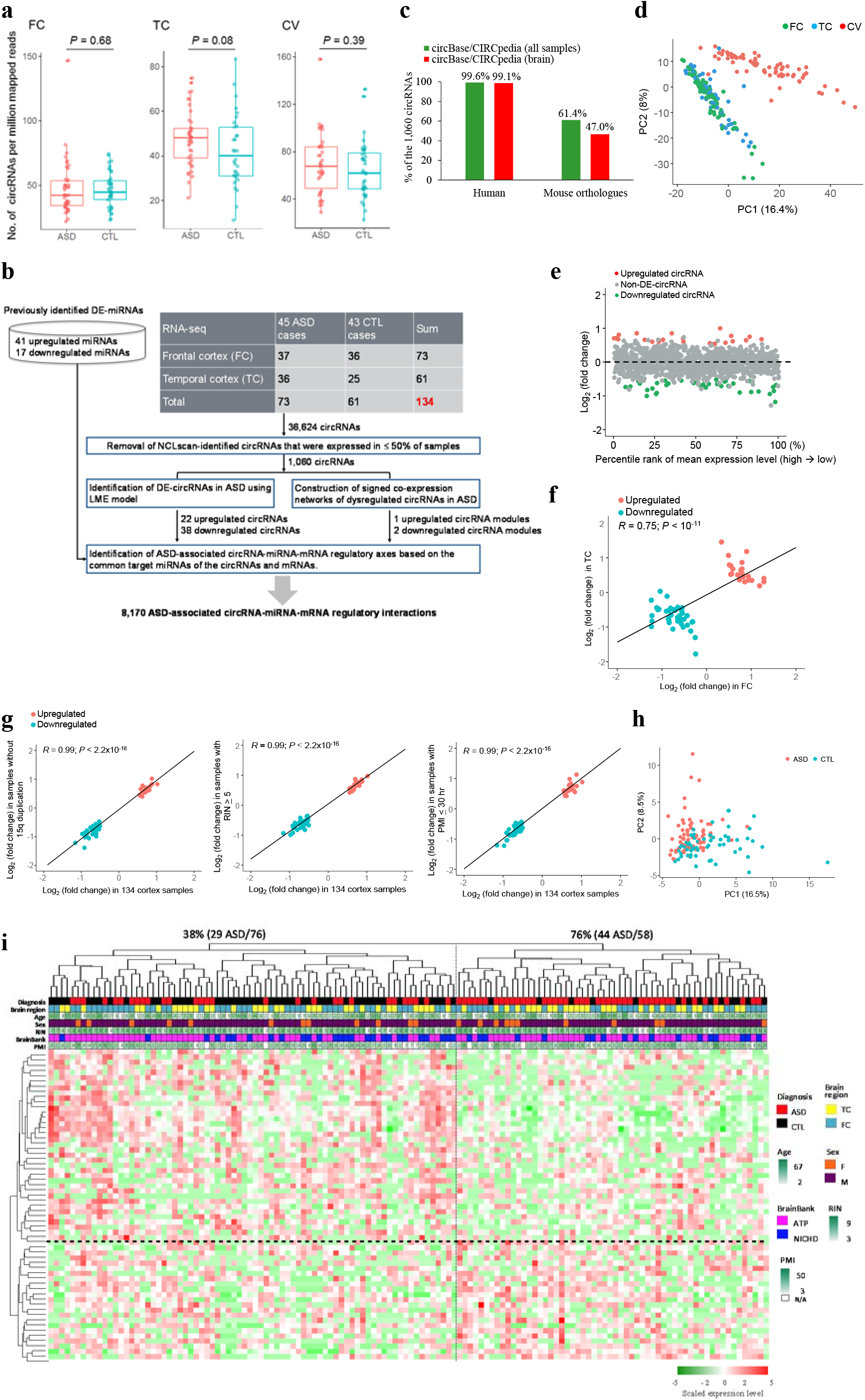
Identification of DE-circRNAs in the ASD cortex samples. (a) Comparison of normalized numbers of circRNAs in ASD and non-ASD control samples from different brain regions (FC, TC, and CV). FC, frontal cortex. TC, temporal cortex. CV, cerebellar vermis. *P* values were determined using two-tailed Wilcoxon rank-sum test. (b) Flowchart of the overall approach. To minimize potentially spurious events, only the circRNAs (1,060 circRNAs) that were detected to be expressed in more 50% of the samples examined are considered for the following analyses. (c) Comparisons of the 1,060 circRNAs and the human/mouse circRNAs collected in well-known databases (i.e., circBase/CIRCpedia). (d) Principal component plots of circRNA expression profiles of the 1,060 circRNAs in samples from FC, TC, and CV. PC1/PC2, the first and second principal components. (e) circRNA expression fold changes (>0 if higher in ASD; <0 if lower in ASD) between ASD and non-ASD control cortex samples, plotted against the percentile rank of mean expression levels of the 1,060 circRNAs across 134 cortex samples used for differential expression analysis. The identified upregulated (22) and downregulated (38) circRNAs in the ASD cortex samples (LME model, *P* < 0.05 and |log_2_(fold change)| ≥ 0.5) are highlighted in red and green, respectively. (f) Comparison of circRNA expression fold changes in the FC and TC samples. The black line represents the regression line between fold changes in the FC and TC for the 60 DE-circRNAs. The Pearson correlation coefficient (*R*) and *P* value are shown. (g) Comparison of circRNA expression fold changes in a small number of samples (left, ASD cases without chromosome 15q11-13 duplication syndrome; middle, ASD cases with RIN ≥ 5; right, ASD cases with PMI ≤ 30h) and all 134 samples combined. The black lines represent the regression lines between fold changes in the corresponding small number of samples and all samples combined for the 60 DE-circRNAs. (h) PCA based on the 60 DE-circRNAs. (i) Dendrogram representing hierarchical clustering of 134 cortex samples based on the identified 60 DE-circRNAs. Information on diagnosis, age, brain bank, PMI, brain region, sex, and RIN is indicated with color bars below the dendrogram according to the legend on the right. Heat map on the bottom represents scaled expression levels (color-coded according to the legend on the bottom) for the 60 DE-circRNAs.

To examine the robustness of the identified DE-circRNAs, we performed resampling analysis with 100 rounds of random sampling of 70% of the samples examined and showed that the fold changes of DE-circRNAs for the resampled and the original sample sets were highly concordant between each other (Pearson’s *R* = 0.95~0.98, all *P* < 2.2×10^−16^; Additional file 4: Figure S1). In addition, we compared the fold changes for a small number of samples (from ASD cases without chromosome 15q11-13 duplication syndrome or with RIN ≥ 5 or post-mortem interval (PMI) ≤ 30h) with those for all samples combined and found a high concordance between each other (all Pearson’s *R* = 0.99, *P* < 2.2×10^−16^; Fig. 1g). These observations revealed that our results were not biased by a small number of samples with removal of chromosome 15q11-13 duplication syndrome, low RIN, or high PMI.

PCA based on the 60 DE-circRNAs revealed that ASD and non-ASD samples could be grouped into separate clusters (Fig. 1h). Moreover, hierarchical clustering showed distinct clustering for the majority of ASD samples. We found two distinct groups for the clustering of the 134 cortex samples, in which 76% (44 out of 58) and 38% (29 out of 76) of cortex samples were ASD samples in the right and left groups, respectively (*P* value < 0.0001 by two-tailed Fisher’s exact test) (Fig. 1i). The confounding factors such as brain region, age, sex, RIN, brain bank, and PMI did not drive the similar clustering (Fig. 1i). Our result thus revealed a shared circRNA dysregulation signature among the majority of ASD samples.

### Construction of signed co-expression networks of circRNA dysregulation in ASD

To probe the correlation between circRNA expression changes and disease status at the system level, we performed the weighted gene co-expression network analysis (WGCNA) to assign individual circRNAs to co-expression modules according to the 1,060 circRNAs identified in the 134 cortex samples (see Methods). As shown in Figure 2a, we identified 14 modules and estimated the module eigengene, i.e., the first principal component (PC1) of the module’s circRNA expression. By assessing the relationship between module eigengene and diagnosis status, three (darkred, violet, and turquoise; Fig. 2b and Additional file 3: Table S3) modules were significantly correlated with ASD status (designated as “DE-modules”), of which the darkred module (including 21 circRNAs) was upregulated and the violet (including 10 circRNAs) and turquoise (including 288 circRNAs) modules were downregulated in ASD samples (Fig. 2c). Of note, the three modules were not correlated with important experimental covariates (age, sex, and brain region) and technique confounders (RIN, PMI, and brain bank) (Fig. 2b).

**Figure 2.**
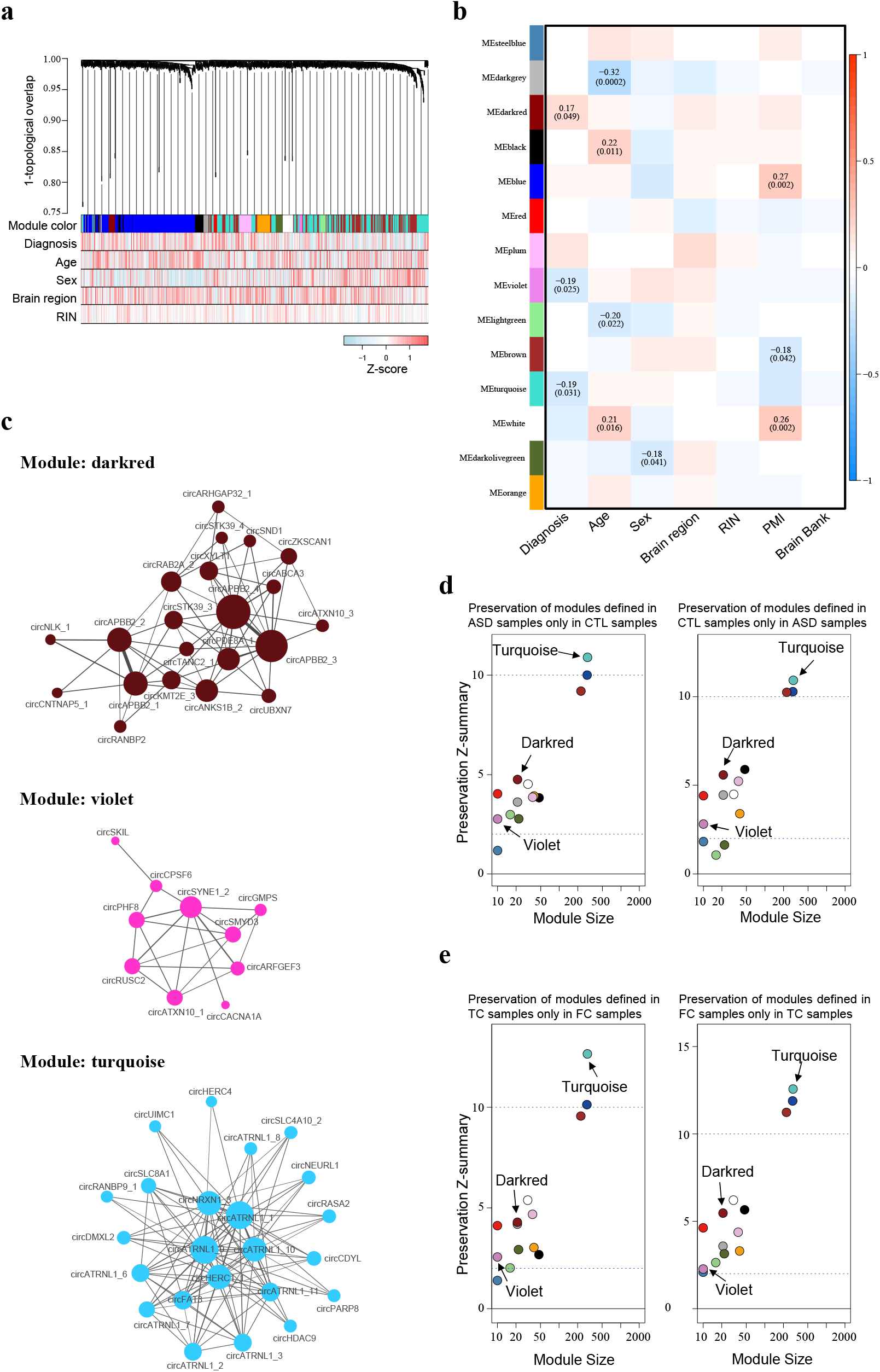
Dysregulation of circRNA coexpression networks in ASD cortex. (a) Dendrogram of circRNA coexpression modules defined in 134 cortex samples. Consensus module color bars shows assignment based on 1,000 rounds of bootstrapping. Diagnosis and potential confounders (age, sex, region, RIN) are treated as numeric variables to calculate their Pearson correlation coefficients with expression level for each circRNA. (b) Pearson’s correlation between distinct covariates and module eigengenes in 134 cortex samples. (c) circRNAs are plotted according their circRNA expression correlations, where the circRNAs in violet and darkred modules are all plotted, but the circRNAs in turquoise module are plotted with only kME 2: 0.5 due to the sizable number of circRNAs. Node size is proportional to connectivity, and edge thickness is proportional to the absolute correlation between two circRNAs. (d, e) Module preservation defined in ASD samples only in CTL samples (d; left), CTL samples only in ASD samples (d; right), TC samples only in FC samples (e; left), or FC samples only in TC samples (e; right).

To test the robustness of the three DE-modules, we then asked if the coexpression structure was similar (1) between ASD and non-ASD control samples and (2) between FC and TC. To this end, we performed module preservation tests [46] (see Methods) to examine preservation of modules defined in ASD samples only in control samples, in control samples only in ASD samples, in TC samples only in FC samples, and in FC samples only in TC samples, respectively. Indeed, we observed that the Z_summary_ scores, which were evaluated for each module to measure the preservation, of all three DE-modules were greater than or equal to two, indicating that the three modules were preserved across ASD and non-ASD control samples (Fig. 2d) and across FC and TC samples (Fig. 2e). Our results thus support the robustness of the three ASD-associated modules.

### Identification of ASD-associated circRNA-miRNA-mRNA regulatory axes

We have identified 60 DE-circRNAs (22 upregulated and 38 downregulated circRNAs) and three DE-modules (one upregulated module including 21 circRNAs and two downregulated modules including 298 circRNAs) in ASD cortex. To further identify the corresponding ASD-associated miRNA-mRNA regulatory axes that were potentially regulated by these ASD-associated circRNAs, we first extracted 58 DE-miRNAs (41 upregulated and 17 downregulated miRNAs) in ASD cortex from the study of Wu et al. [15] These ASD-affected miRNAs were derived from 95 human cortex samples (47 samples from 28 ASD cases and 48 samples from 28 non-ASD controls), of which 73 samples overlapped with the samples examined in this study. Since circRNAs often act as a miRNA sponge and the function can be enhanced through the long half-lives of circRNAs, we directly assessed the role of circRNA dysregulation in alterations of the ASD-affected miRNAs. To achieve this, we searched for the target sites of the 41 upregulated (or 17 downregulated) miRNAs in the circle sequences of the 38 downregulated circRNAs and 298 circRNAs defined in two downregulated circRNA modules (or 22 upregulated circRNAs and 21 circRNAs defined in one upregulated circRNA module) (Additional file 4: Figure S2; see Methods). After that, we determined the ASD-associated circRNA-miRNA-mRNA interactions by integrating the circRNA-miRNA interactions with the miRNA-mRNA interactions (Additional file 5: Table S4) according to the common miRNA target sites of the circRNAs and mRNAs (see Methods). We then calculated the correlations between circRNA and miRNA expression, between miRNA and mRNA expression, and between circRNA and mRNA expression based on the same set of cortex samples (i.e., 73 samples). As illustrated in Figure 3a, only the circRNA-miRNA-mRNA interactions were considered if they simultaneously satisfied the following rules (see also Methods): (1) the circRNA-miRNA axes should be upregulated circRNA-downregulated miRNA or downregulated circRNA-upregulated miRNA axes; (2) both the circRNA-miRNA and miRNA-mRNA interactions should exhibit a significantly negative correlation of expression profile between circRNAs and the corresponding predicted regulated miRNAs and between miRNAs and target mRNAs, respectively; (3) the circRNA expression should be positively correlated with the corresponding mRNA expression; and (4) the Fisher’s combined *P* values [47] of the above three independent Spearman’s correlation tests should be less than 0.05. These criteria led to a total of 8,170 (including 356 upregulated and 7,814 downregulated circRNA-involved interactions) ASD-associated circRNA-miRNA-mRNA axes, which involved 2,302 target genes (Table 2 and Additional file 6: Table S5).

**Table 2.** Brain tissue samples used in this studies.

|  |  |
| --- | --- |
| No. of interactions | 8,170 (356 upregulated circRNA-involved and 7,814 downregulated circRNA-involved interactions) |
| No. of DE-circRNAs | 46 (3 upregulated and 43 downregulated circRNAs) |
| No. of DE-miRNAs | 30 (26 upregulated and 4 downregulated miRNAs) |
| No. of target genes | 2,302 |

Note. DE, differentially expressed.

**Figure 3.**
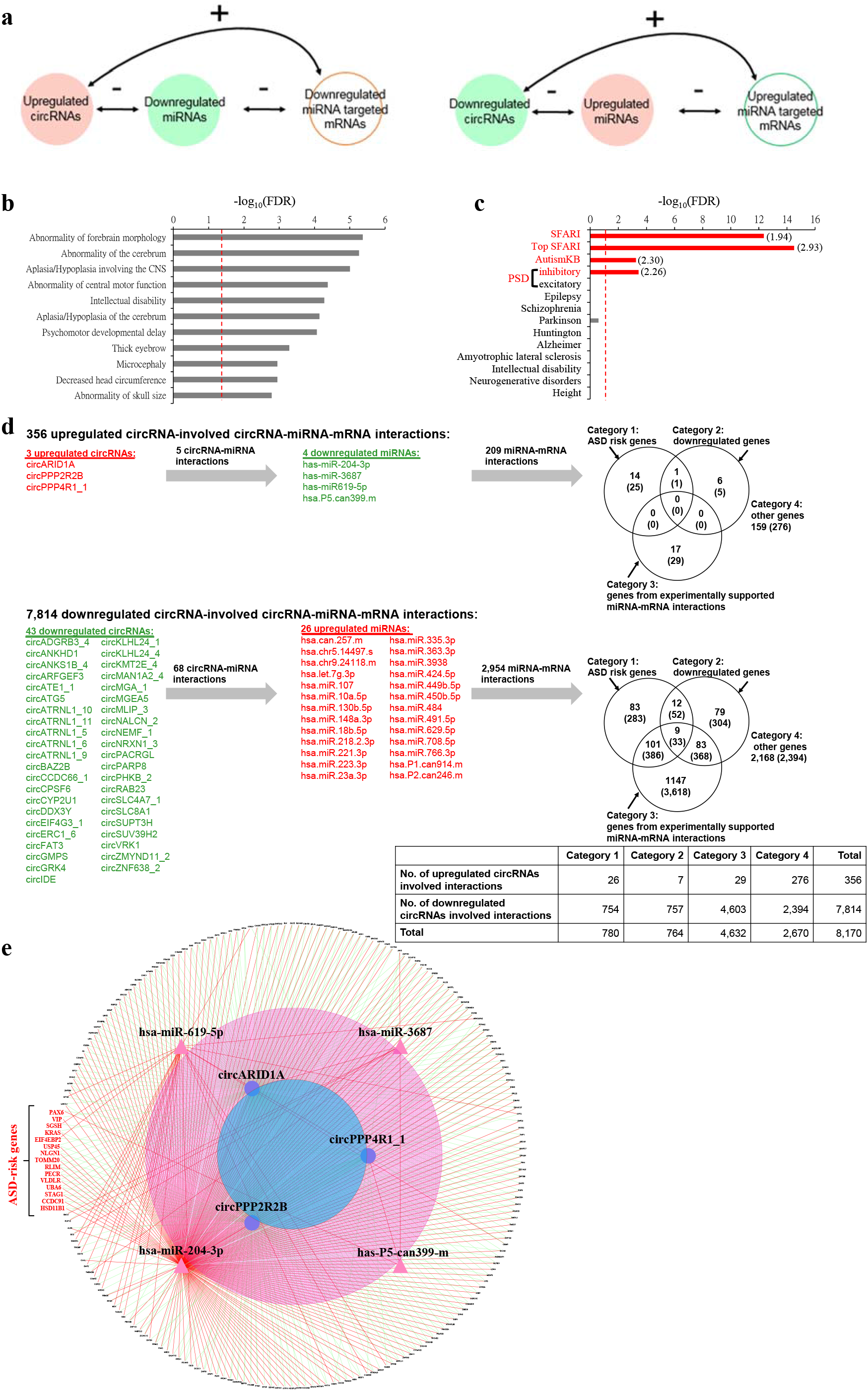
Identification of ASD-associated circRNA-miRNA-mRNA regulatory interactions based on the 73 cortex samples overlapped with the samples examined in a previous miRNA study[15]. (a) Schematic diagram representing the ASD-associated circRNA-miRNA-mRNA axes that satisfied our criteria (see the text). “-” represents the negative correlation between the expression of circRNAs and miRNAs and between the expression of miRNAs and mRNAs. “+” represents the positive correlation between the expression of circRNAs and mRNAs. The correlations are performed by Spearman’s correlation tests. (b,c) Enrichment analyses of phenotype ontology (b) and ASD risk genes (c) among the target genes (2,302 genes) of the identified ASD-associated circRNA-miRNA-mRNA axes. The red dashed lines represent FDR Bonferroni-corrected *P* values for enrichments with FDR < 0.05. For (c), the enrichment odds ratios with FDR < 0.05 are provided in parentheses. Top SFARI genes represent the genes with top ASD genetic risk factors from the SFARI database (score = 1-3 and syndromic). (d) The four categories of upregulated (top) and downregulated (bottom) circRNA-involved ASD-associated circRNA-miRNA-mRNA interactions. The Venn diagrams represent the overlap between the four categories of interactions, on which the numbers between brackets show the numbers of target genes. The numbers of the identified circRNA-miRNA-mRNA interactions are shown in parentheses. A summary table representing the total number of interactions for each category is also provided (bottom right). (e) The 356 upregulated circRNA-involved circRNA-miRNA-mRNA interactions plotted by the Cytoscape package.

To explore the relationship between ASD and the 2,302 target genes of these ASD-associated circRNA-miRNA-mRNA interactions, we first assessed shared phenotypes among the target genes by enrichment for Human Phenotype Ontology terms (the ToppFunn module of ToppGene Suite software [48]). We found that the target genes were significantly enriched in the phenotype ontology terms of aplasia/hypoplasia involving the central nervous system (CNS) and abnormality of forebrain morphology, the cerebrum, the cerebral subcortex, and skull size (Fig. 3b), reflecting the brain morphometry differences between ASD patients and healthy individuals [49]. Regarding the 2,302 target genes, we further examined enrichment for ASD risk genes from Simons Foundation Autism Research Institutive (SFARI) AutDB [3] and AutismKB [5] databases (SFARI and AutismKB genes), which have been previously implicated in ASD through genetic syndromes, candidate gene studies, common variant association, and structural variation. Intriguingly, the target genes showed significant enrichment for both SFARI and AutismKB genes (particularly for top SFARI genes with score = 1-3 and syndromic; all FDR < 0.05 by two-tailed Fisher’s exact test), but not for genes implicated in monogenetic forms of schizophrenia, Alzheimer’s disease, intellectual disability, and other brain disorders (Fig. 3c). These results suggest that targets of the identified circRNA-miRNA-mRNA axes are enriched for genes causally connected with autism, but less so far genes connected with other brain disorders. In addition, we found that these target genes were significantly enriched (FDR < 0.05) for genes encoding inhibitory postsynaptic density (PSD) proteins, but not for those encoding excitatory PSD ones (Fig. 3c). This result is consistent with a previous observation that ASD-derived organoids exhibit overproduction of inhibitory neurons [50].

According to the target genes previously implicated in ASD or the experimental evidence of miRNA-mRNA binding, we further classified the identified circRNA-miRNA-mRNA interactions into four categories as follows (Fig. 3d).

*Category 1*: The target genes have been previously reported to be ASD risk genes (i.e., SFARI or AutismKB genes).
*Category 2*: The target genes were reported to be DE-genes in ASD [12] based on the samples overlapped with the samples examined in this study. The interactions should be either upregulated circRNA-downregulated miRNA-upregulated mRNA or downregulated circRNA-upregulated miRNA-downregulated mRNA interactions.
*Category 3*: The miRNA-mRNA binding has been experimentally validated (see Methods).
*Category 4*: Other.

The ASD-affected circRNAs (DE-circRNAs and circRNAs in the DE-modules) of the interactions may provide an upstream regulation for the ASD-associated miRNA-mRNA axes and thereby potentially contribute to ASD susceptibility (see also Fig. 3e and Additional file 4: Figure S3). Particularly, Category 1 interactions, which included 188 target ASD risk genes (Additional file 6: Table S5), may play an important regulatory role in ASD brain.

### Experimental validation of the negative regulation of downregulated miRNAs by upregulated circRNAs in human neuronal cell lines

We proceeded to assess the role of circRNA dysregulation in ASD-associated miRNA level alterations. Of the identified circRNA-miRNA-mRNA interactions (Fig. 3d), we selected a DE-circRNA (the upregulated circARID1A) that was predicted to have the greatest number of target sites of miRNAs that have been identified to be differentially expressed in the same ASD samples used in this study [15]. circARID1A was predicted to harbor the largest number of target sites of one single downregulated miRNA (seven miR-204-3p target sites; Fig. 4a and Additional file 4: Figure S2). The circRNA junction of circARID1A has not yet been experimentally confirmed previously. The role of circARID1A and its regulatory interaction with miR-204-3p are unknown, and has not yet been studied experimentally. We first confirmed the non-co-linear (NCL) junction (or the backspliced junction) of circARID1A. Since comparisons of different RTase products have been demonstrated to effectively detect RT-based artificial NCL junctions [26, 51–54], we employed reverse transcription polymerase chain reaction (RT-PCR) using Avian Myeloblastosis Virus (AMV)- and Moloney Murine Leukemia Virus (MMLV)-derived RTase in parallel experiments in the examined cell lines/tissues, followed by Sanger sequencing of the RT-PCR amplicons to validate the NCL junction (Fig. 4a). Our result showed that the NCL junction of circARID1A was RTase-independent (Fig. 4a), supporting that the NCL junction was unlikely to be generated from an RT-based artifact. We further treated total RNA from the examined cell lines/tissues with RNase R and showed the RNase R-resistance of the NCL junction, supporting the existence of the circRNA event (Fig. 4b). Intriguingly, we observed that circARID1A was commonly expressed across vertebrate (from primates to chicken) brains, indicating the evolutionary significance of circARID1A (Fig. 4c). We further examined the expression profiles of circARID1A and its corresponding co-linear mRNA counterpart in various human tissues and found that these two isoforms exhibited different expression patterns (Fig. 4d). While circARID1A was particularly enriched in the brain, its corresponding co-linear mRNA counterpart was not (Fig. 4d). Importantly, regarding the relative expression of these two isoforms in the brain, circARID1A was significantly more abundant than its co-linear form; in contrast, circARID1A was expressed at a relatively low level as compared to its co-linear form in the non-brain tissues (Fig. 4e). We also found that circARID1A was widely expressed in 24 human brain regions (Additional file 4: Figure S4). Together, these results suggested that circARID1A may play important biological roles in human brain.

**Figure 4.**
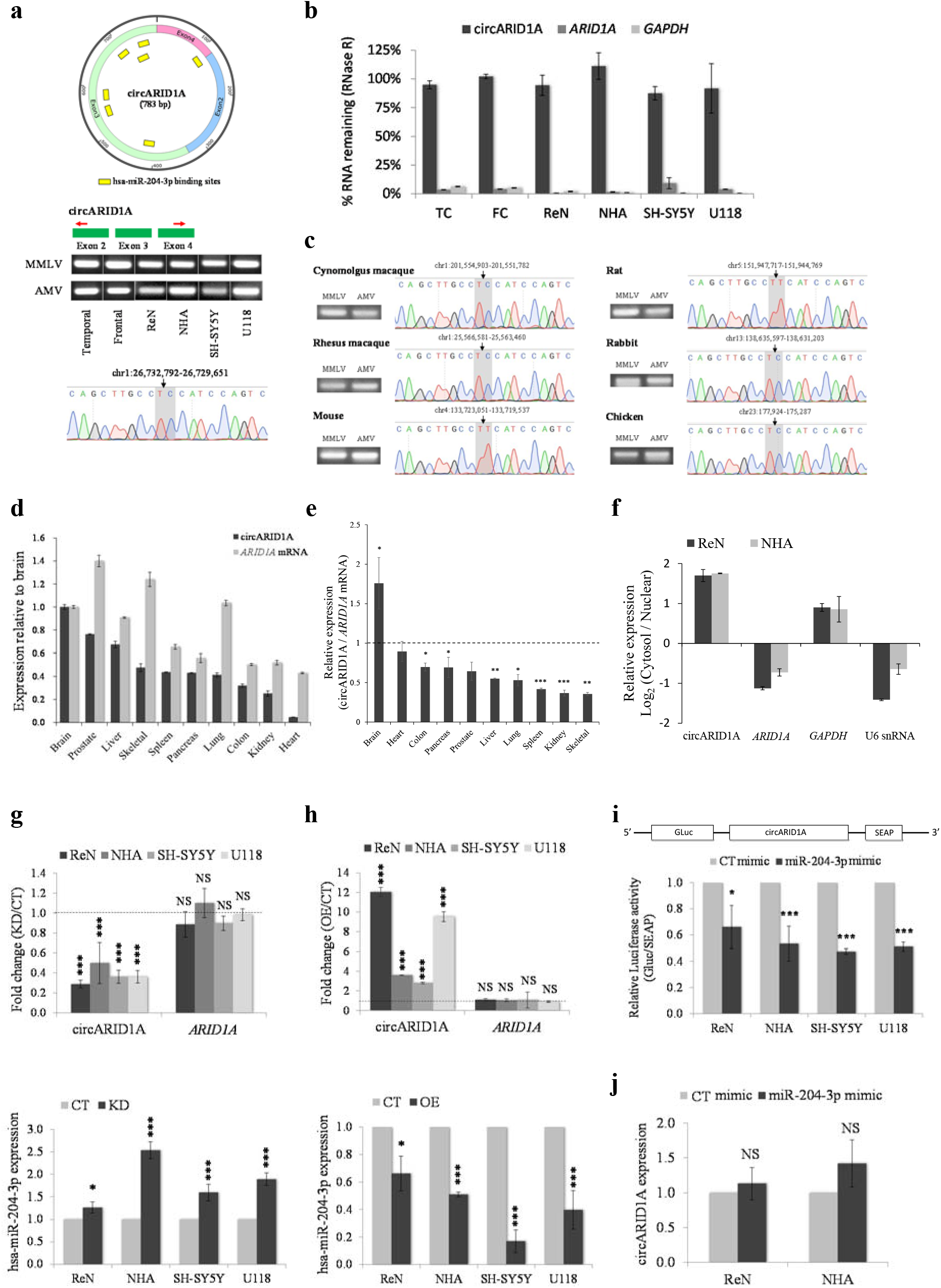
Experimental validation of the backspliced junction of an upregulated circRNA (circARID1A) and the corresponding circRNA-miRNA regulatory axis. (a) Validation of the NCL junction of circARID1A. Top and middle: the schematic diagrams displaying seven predicted target sites of one single downregulated miRNA (miR-204-3p) on the circle of circARID1A (top) and the designed divergent primers around the NCL junction of circARID1A (middle). Bottom: comparisons of two different RTase products (MMLV- and AMV-derived products) of circARID1A in TC/FC samples and four types of neuronal cell lines (hNPC (or ReN), NHA, SH-SY5Y, and U118), followed by Sanger sequencing the RT-PCR amplicons for the NCL event in the TC. ReN, ReNcell VM. (b) Experimental validation of the circRNA (or backspliced) junction of circARID1A. The figure shows the expression fold changes (as determined by qRT-PCR) for circARID1A, *ARID1A* mRNA, and *GAPDH* (negative control) in the indicated tissues/cell lines before and after RNase R treatment. (c) Experimental examination of the evolutionary conservation of circARID1A across the brains of vertebrate species from primates to chicken. Comparison of MMLV- and AMV-derived-RTase products (left) and the corresponding sequence chromatograms (right) for the circARID1A event in the brains of the indicated six species are shown. (d) Comparison of the expression profiles of circARID1A and its corresponding co-linear mRNA counterpart in 10 normal human tissues. The expression levels of brain are used to normalize the relative expression values of the other tissues. (e) The relative expression of circARID1A and its corresponding co-linear mRNA counterpart in 10 normal human tissues. (f) qRT-PCR analysis of the cytoplasmic to nuclear expression ratios for circARID1A and *ARID1A* mRNA. *GAPDH* and U6 snRNA are examined as controls. (g,h) qRT-PCR analyses of the correlations between the expression of circARID1A and miR-204-3p after circARID1A knockdown (g) or overexpression (h) in various human neuronal cell lines. The top panels of (g) and (h) represent that circARID1A knockdown (g) or overexpression (h) did not significantly affect the *ARID1A* mRNA expression. CT, control. KD, knockdown. OE, overexpression. (i) Luciferase reporter assay for the luciferase activity of Gaussia luciferase (GLuc)-circARID1A in ReN, NHA, SH-SY5Y, and U118 cells transfected with miR-204-3p mimics to validate the binding between circARID1A and miR-204-3p. The entire circle sequence of circARID1A was cloned into the downstream region of the GLuc gene (i.e., GLuc-circARID1A; top). The luciferase activity of GLuc was normalized with secreted alkaline phosphatase (SEAP). (j) qRT-PCR analysis of the expression level of circARID1A in ReN and NHA cells after transfection with miR-204-3p mimics. All the qRT-PCR data are the means ± SEM of three experiments. *P* values are determined using two-tailed *t* test. Significance: \**P* value < 0.05 and \*\*\**P* value < 0.001. NS, not significant.

To test if circARID1A regulates miR-204-3p activities, we first confirmed that circARID1A was indeed predominantly expressed in the cytoplasm in both NHAs and hNPCs (e.g., ReNcell VM; designated “ReN”) (Fig. 4f). We then examined the correlation between expression of circARID1A and miR-204-3p in different types of neuronal cell lines. To this end, we disrupted circARID1A expression and overexpressed circARID1A, respectively (see Methods). We found that circARID1A knockdown (Fig. 4g, top) and overexpression (Fig. 4h, top) did not significantly affect the expression of its corresponding co-linear mRNA counterpart (*ARID1A*). However, miR-204-3p expression indeed significantly increased and decreased after circARID1A knockdown (Fig. 4g, bottom) and overexpression (Fig. 4h, bottom), respectively. To further examine if miR-204-3p can bind to circARID1A, we performed luciferase reporter assay by co-transfecting the miRNA mimic with the luciferase reporters into NHAs and hNPCs. We showed that miR-204-3p significantly reduced the luciferase reporter activities (Fig. 4i), confirming the binding between circARID1A and miR-204-3p. We also showed that miR-204-3p overexpression did not significantly affect circARID1A expression (Fig. 4j). These results thus confirmed the negative regulation between circARID1A and miR-204-3p and the regulatory role of circARID1A as a miR-204-3p sponge.

### Regulation of ASD risk genes via the identified circRNA-miRNA axis

Our results showed that circARID1A can indeed function as a sponge to neutralize the activity of miR-204-3p (Figs. 4g-4j). Of the identified circRNA-miRNA-mRNA regulatory interactions, there were 171 target genes (including 12 ASD risk genes) in the circARID1A-miR-204-3p-involved interactions (Fig. 5a and Additional file 4: Figure S5). We next experimentally tested the predicted upregulation and downregulation of target genes by circARID1A and miR-204-3p in ASD via *in vitro* perturbation of circARID1A and miR-204-3p in hNPCs, respectively. To this end, we overexpressed either circARID1A or miR-204-3p in hNPCs and examined the fold changes of the target gene expression by microarray analysis (Additional file 7: Table S6). If the target genes are regulated by the circARID1A-miR-204-3p axis, there should be a negative correlation between the mRNA fold changes after circARID1A overexpression and those after miR-204-3p overexpression. Indeed, we found that overexpression of circARID1A significantly upregulated the target genes as compared to those with miR-204-3p overexpression (*P* < 10^−4^ by Kolmogorov-Smirnov test; Fig. 5b). In addition, mRNA fold changes after circARID1A overexpression were negatively correlated with those after miR-204-3p overexpression (Pearson’s *R* = −0.7, *P* < 10^−5^; Fig. 5c). For the 12 ASD risk genes, circARID1A overexpression led a pronounced increase, whereas the reverse was observed for miR-204-3p overexpression (*P* < 0.05 by one-tailed paired *t* test; Fig. 5d). These results were consistent with our bioinformatics predictions.

**Figure 5.**
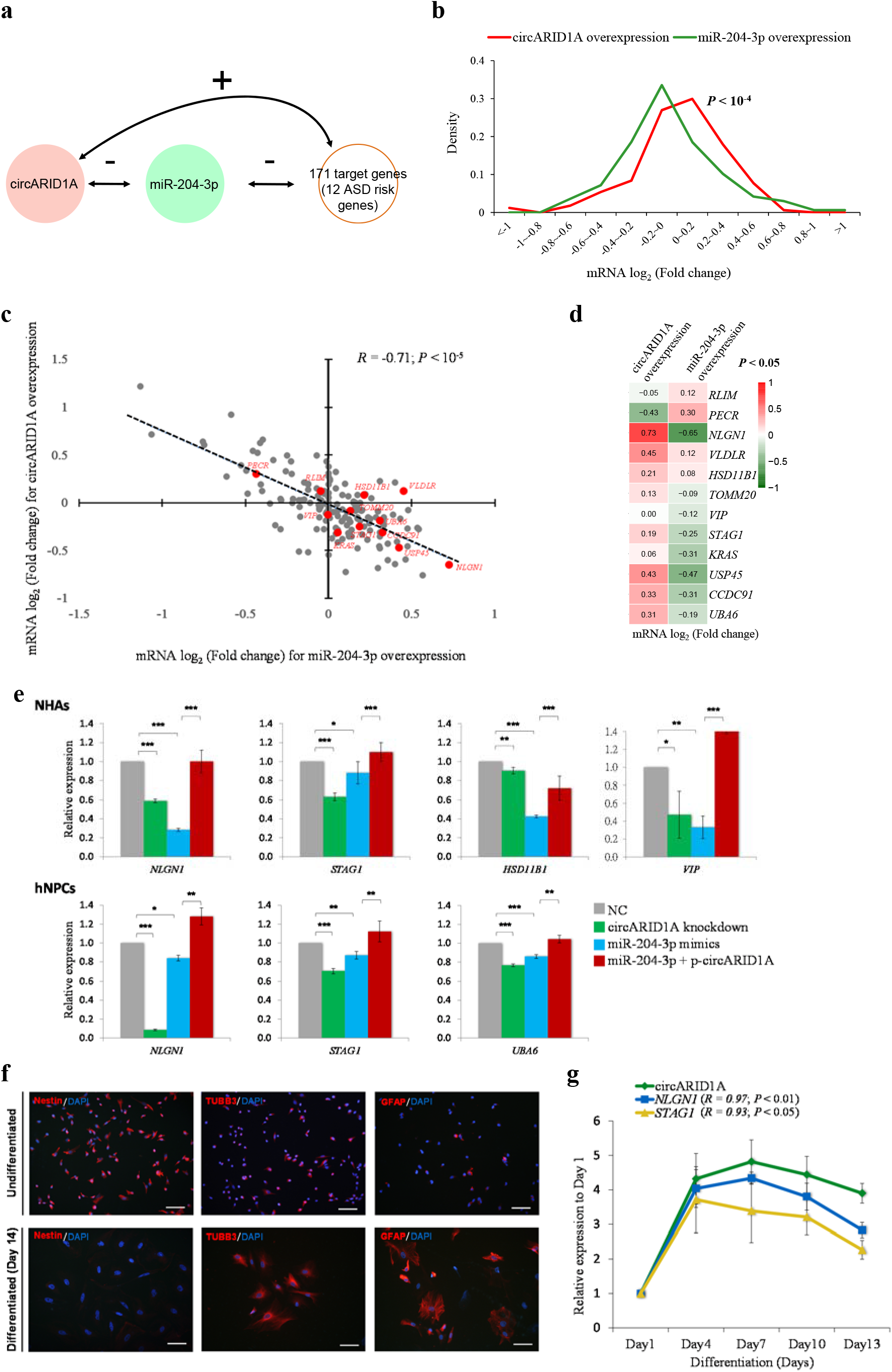
Experimental validation of the circARID1A regulatory role of miR-204-3p sponges and the corresponding circRNA-miRNA-mRNA regulatory interactions. (a) Schematic diagram representing the circARID1A-miR-204-3p regulatory axis and its regulated miRNA targets (171 genes, including 12 ASD risk genes). (b-d) Microarray analysis of the target mRNA log_2_(fold change) in response to overexpression of circARID1A and miR-204-3p. (b) Distribution of the target mRNA log_2_(fold change) in response to overexpression of circARID1A and miR-204-3p. *P* value is determined using Kolmogorov-Smirnov test. (c) A negative correlation between mRNA fold changes after circARID1A overexpression and miR-204-3p overexpression. The red dots represent the 12 ASD risk genes. The Pearson’s correlation coefficient (*R*) and *P* value are shown. (d) Heat map of the 12 ASD risk mRNA log_2_(fold change) in response to overexpression of circARID1A and miR-204-3p. *P* value is determined using one-tailed paired *t* test. (e) qRT-PCR analyses of ASD risk gene expression in NHAs (top) and hNPCs (ReNcell VM or ReN) (bottom) cells after circARID1A knockdown, miR-204-3p mimics, and miR-204-3p mimics with circARID1A expression vector (p-circARID1A), respectively. *P* values are determined using two-tailed *t* test. Significance: \**P* value < 0.05, \*\**P* value < 0.01, and \*\*\**P* value < 0.001. (f) Imaging of immunostained ReN cells after two weeks of differentiation. Immunostaining shows undifferentiated (top) and differentiated (bottom) ReN cells with the neural stem cell marker nestin (left), the neuronal marker βIII-tubulin (middle), and the mature glial cell marker GFAP (right). Nuclei are stained with DAPI (blue). Scale bar = 100 μm. GFAP, glial fibrillary acidic protein. (g) Relative expression of circARID1A and two ASD risk genes (*NLGN1* and *STAG1*) during hNPC differentiation. The Pearson correlation coefficients (*R*) between the expression of circARID1A and the two ASD risk genes and *P* values are shown in in parentheses. All the qRT-PCR data are the means ± SEM of three experiments.

Next, we experimentally examined the regulation of the circARID1A-miR-204-3p axis on several ASD risk genes NHAs and hNPCs. In NHAs, four SFARI genes (*NLGN1*, *STAG1*, *HSD11B1*, and *VIP*) were significantly downregulated by the knockdown of circARID1A and the overexpression of miR-204-3p, respectively (Fig. 5e). The miR-204-3p-mediated repression of the four SFARI genes was rescued by ectopic expression of circARID1A. Similarly, in hNPCs, we found that three SFARI genes (*NLGN1*, *STAG1*, and *UBA6*) were significantly downregulated by circARID1A knockdown and miR-204-3p mimics, and the miR-204-3p-mediated repression of the three genes was rescued by ectopic expression of circARID1A (Fig. 5e). Furthermore, we differentiated hNPCs into different neural cell types (Fig. 5f) and found that circARID1A expression was significantly positively correlated with the expression of *NLGN1* and *STAG1* during hNPC differentiation (Fig. 5g). Taken together, these results suggest that circARID1A could regulate some ASD risk genes by directly sponging miR-204-3p.

## Discussion

With a large brain sample size of both small RNA-seq data and RNA-seq data from total RNAs (rRNA-depleted RNAs without polyA-selection) of ASD cases and controls, the Synapse database [12] brought an unprecedented opportunity for us to conduct the first study, to the best of our knowledge, for systematically investigating circRNA dysregulation in ASD and the corresponding ASD-associated circRNA-miRNA-mRNA regulatory axes. Our genome-wide integrative analysis thus provided new insights into the role of circRNAs in ASD pathophysiology. We identified 60 DE-circRNAs and three DE-modules of circRNAs in ASD cortex, and showed a shared circRNA dysregulation signature among the majority of ASD samples. Regarding the identified ASD-associated circRNAs and the previously identified DE-miRNAs [15] derived from the same cortex samples used in this study, we identified the ASD-associated circRNA-miRNA regulatory axes and thus constructed 8,170 ASD-associated circRNA-miRNA-mRNA interactions. Importantly, for the target genes (i.e., 2,302 genes) of the 8,170 interactions, we observed significant enrichment for ASD risk genes (top ASD risk genes particularly), but not for genes implicated in monogenetic forms of other brain disorders (such as epilepsy, which is often a comorbidity of ASD) (Fig. 3c). On the basis of a most recently released dataset of high-confidence ASD genetic risk genes [55] (102 genes; see Methods), the targets also exhibited significant enrichment for the high-confidence ASD genetic risk genes (*P* < 10^−4^ by two-tailed Fisher’s exact test; Additional file 7: Figure S6). Moreover, we observed that target genes of the ASD-associated circRNA-miRNA-mRNA axes were significantly enriched in genes encoding inhibitory PSD proteins, but not in those encoding excitatory PSD ones (Fig. 3c). This result reflects the previous reports that there is an excitatory-inhibitory neuron imbalance in ASD [56, 57] and that inhibitory neurons are overproduced in ASD patient-derived organoids [50]. These results suggest that the identified ASD-associated circRNA-miRNA axes may serve as an alternative pathway for gene-disrupting mutations to perturb key transcript levels and thereby contribute to ASD susceptibility. Our experimental validation showed the negative correlation between the expression changes of ASD-affected circRNAs and miRNAs (Figs. 4g and 4h) and the regulation of the target ASD risk genes by the corresponding circRNA-miRNA axis (Fig. 5e). Our results thus suggest that circRNAs can function as efficient miRNA sponges for upstream regulation of the corresponding ASD risk genes.

Regarding the overall circRNA expression profiles, our PCA result represented that two types of cortex samples (FC and TC) were clustered together but cortex and CV samples were grouped into separate clusters (Fig. 1d). This result is similar to a new separate study for circRNA expression analysis (Gokoolparsadh et al., preprint) [58] and some previous observations for mRNA [11, 45] and miRNA [15] expression profiles. This also reflects the phenomenon that transcriptomic profiles are commonly clustered according to broad brain region [45]. Meanwhile, like the pattern observed by Gokoolparsadh et al. [58], our result (Additional file 4: Figure S7) also revealed that ASD and non-ASD samples did not show distinct clustering based on the overall circRNA expression profiles of all circRNAs identified from the cortex samples[58]. However, on the basis of the DE-circRNAs identified in this study, the two types of samples (ASD and non-ASD samples) can be grouped into two separate clusters by both PCA (Fig. 1h) and hierarchical clustering (Fig. 1i) analyses. We showed that the fold changes for the DE-circRNAs were concordant between the FC and TC (Fig. 1f) and were not biased by a small number of samples with removal of chromosome 15q11-13 duplication syndrome, low RIN, or high PMI (Fig. 1g), supporting the robustness of the identified DE-circRNAs. We thus successfully provided a shared circRNA dysregulation signature among the majority of ASD samples.

Of the 8,170 identified ASD-associated circRNA-miRNA-mRNA interactions, the circRNA-miRNA axes should be either upregulated circRNA-downregulated miRNA axes or downregulated circRNA-upregulated miRNA axes (Fig. 3a). To benefit the further studies for the identified circRNA-miRNA-mRNA regulatory interactions in ASD susceptibility, we divided these 8,170 circRNA-miRNA-mRNA interactions into four categories according to the target genes previously implicated in ASD or the experimental evidence of miRNA-mRNA binding (Fig. 3d). Particularly, Category 1 interactions (780 interactions; Fig. 3d), in which the target genes have been previously implicated in ASD, provided a useful set of upstream regulations (circRNA-miRNA axes) for the ASD risk genes. As ASD risk genes, which have been implicated through ASD rare or *de novo* variations, are predicted to disrupt protein function, downregulated circRNAs may result in miRNA upregulation in ASD and thereby contribute to ASD risk genes. On the other hand, upregulated circRNAs, which may act as a role of miRNA sponges, may result in miRNA downregulation in ASD. A previous study [15] suggested that downregulated miRNAs may play a compensatory or adaptive role for causing upregulation of ASD risk genes. In addition to the ASD risk gene-involved interactions (i.e., Category 1), we emphasized that all the identified circRNA-miRNA-mRNA regulatory axes were derived from aberrant circRNA and miRNA expression in ASD and the correlations between circRNA, miRNA, and mRNA expression through the same cortex samples. The target genes of Categories 2-4 interactions, which were identified to be regulated by upstream ASD-affected circRNA-miRNA axes, may be valuable candidates for further studies in idiopathic ASD.

In this study, we not only provided a systems-level view of landscape of the circRNA expression in ASD cortex samples and the corresponding ASD-associated circRNA-miRNA-mRNA axes, but also experimentally characterized the targets of an upregulated circRNA-downregulated mRNA axis (i.e., the circARID1A-miR-204-3p axis) in NHAs and hNPCs. We confirmed that the expression of five SFARI genes (*NLGN1*, *STAG1*, *HSD11B1*, *VIP*, and *UBA6*) were regulated by circARID1A via sponging miR-204-3p in NHAs or hNPCs (Fig. 5e). The expression of both *NLGN1* and *STAG1* exhibited a significantly positive correlation with the circARID1A expression during hNPC differentiation (Fig. 5g). Of note, *NLGN1* has been reported to play an important role in a variety of activity-dependent response [59] and memory formation [60–62]. Knockout of *NLGN1* in mice could cause increased repetitive behavior [62]; and loss or overexpression of *NLGN1* could impair spatial memory in transgenic mouse models. Alteration of *NLGN1* expression in specific excitatory and inhibitory neuronal subpopulations can affect the dynamic processes of memory consolidation and strengthening [60]. Since excitatory-inhibitory neuron imbalances often accompany neuropsychiatric disorders, the dysregulation of *NLGN1* may underlie the disorders. Therefore, the identified circARID1A-miR-204-3p axis, which regulates *NLGN1* expression, may provide a useful circuitry/molecular mechanism of excitation and inhibition underlying long-term memory consolidation and strengthening for further developing potential therapeutic strategies to address these neuropsychiatric disorders, including ASD. In addition, *VIP* and *UBA6* have been demonstrated to play an essential regulatory role during rodent embryonic development [63, 64]. *VIP* is known as a regulator of embryogenesis of neural tube closure; interference with *VIP* can result in permanent effects on adult social behavior [64]. It was shown that *UBA6* brain-specific knockout mice exhibited social impairment and reduced vocalizations, representing a valid ASD mouse model [65]. As a regulator of multiple ASD-associated genes, the circARID1A-miR-204-3p axis would be a valuable candidate for further ASD study.

It is noteworthy that circARID1A was validated to be widely expressed in the brain of multiple vertebrate species from human to chicken (Fig. 4c), suggesting the evolutionary importance of this circRNA across vertebrate species. This observation also implies a possibility that future studies may further examine whether circARID1A serves as an early pathogenic feature during neurodevelopmental processes by altering circARID1A expression in appropriate transgenic animal models such as mice. In addition, we observed circARID1A was widely expressed in multiple brain regions (Additional file 4: Figure S4). This raises an interesting question of whether circARID1A dysregulation could occur in multiple cell types or cell composition in ASD brains. Two recent studies have comprehensively investigated gene dysregulation in ASD [66] and Alzheimer’s disease (AD) [67] in a cell type-specific manner by single-nucleus RNA sequencing of cortical tissue from patients with ASD or AD, respectively. This issue may be addressed if the approach of single-cell sequencing from total RNA samples without polyA-selection is available in the future.

Taken together, our study has provided a framework for assessing the functional involvement of circRNA in ASD and the corresponding ASD-associated circRNA-miRNA-mRNA regulatory axes. By integrating ASD candidate gene sets and circRNA, miRNA, and mRNA dysregulation data derived from the same ASD cortex samples, we have provided multiple lines of evidence for the functional role of ASD for circRNA dysregulation and a rich set of ASD-associated circRNA candidates and the corresponding circRNA-miRNA-mRNA axes, particularly those involving ASD risk genes, for further investigation in ASD pathophysiology.

## Methods

### Identification of circRNAs in the cortex samples

The rRNA-depleted RNA sequencing data of human brain was obtained by the request from Synapse (http://www.synapse.org) under the accession number syn4587609, which included 236 post-mortem samples of frontal cortex (FC), temporal cortex (TC), and cerebellar vermis (CV) from 48 ASD-affected individuals and 49 non-ASD-affected controls (Table 1). We first applied NCLscan to circRNA identification in these ASD and non-ASD control samples on the basis of human reference genome (GRCh38) and the Ensembl annotation (version 90). The expression levels of circRNAs were calculated using the number of supporting circRNA junction reads per million uniquely mapped reads (*RPM*) [68]. For accuracy, a sample was not considered in the following analysis if the number of the identified circRNAs of this sample was one standard deviation below the mean of the sample set. Therefore, 10 CV and 24 cortex (12 FC and 12 TC) samples were removed in this study (i.e., 202 samples (73 FC, 61 TC, and 68 CV samples) were retained). There were 53,427 NCLscan-identified circRNAs in the 202 samples (Additional file 2: Table S2). Since several independent transcriptomics studies of ASD have indicated that human cortex has been implicated in ASD pathophysiology [41, 42] and changes in transcriptomic profiles were stronger in the cortex than in the cerebellum [12], our following analysis focused on the 134 samples of prefrontal and temporal cortex (73 FC and 61 TC samples) (Fig. 1b). Of the 134 cortex samples, 36,624 circRNAs were identified. To minimize potentially spurious events, we only considered the circRNAs that were detected in more 50% of the 134 samples examined. Thus, a total of 1,060 circRNAs were used in the following analyses. Regarding these 1,060 circRNAs, we performed principal component analysis (PCA) and showed that circRNA expression profiles were very similar between two types of cortex regions but were quite distinct in the cerebellum (Fig. 1b). The previously identified human and mouse circRNAs were downloaded from two publicly accessible datasets: circBase (www.circbase.org)[43] and CIRCpedia v2 (www.picb.ac/rnomics/circpedia/) [44]. Since the genomic coordinates of human circRNAs in circBase were defined on the hg19 assembly, we used the liftOver tool [69] to obtain the genomic coordinates of circRNAs on the GRCh38 assembly. Regarding the mouse circRNAs collected in circBase (mm9) and CIRCpedia v2 (mm10), we also performed the liftOver tool to obtain the corresponding orthologous mouse coordinates of circRNAs on the GRCh38 assembly.

### Identification of differentially expressed circRNAs in ASD

We then used the nlme program in the R package to perform a linear mixed effects (LME) model and detect DE-circRNAs between ASD and control samples (diagnosis) with controlling for potential confounding factors including sex, age, brain region (FC or TC), RNA quality (RNA integrity number; RIN), host gene expression, sequencing batch, and brain bank batch as follows:

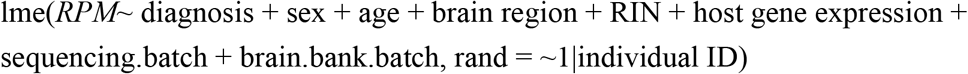

Here, each individual sample was set as a random effect; the other factors were set as fixed effects. Significant results of the diagnosis factor were reported at *P* < 0.05. Of note, the expression levels of the host genes were measured by log_2_ normalized FPKM (which accounts for gene read counts, GC content, gene length, and library size) using the cqn program in the R package [70]. The read counts of the host genes were calculated by the STAR aligner [71], followed by the RSEM tool [72]. We thereby identified 60 DE-circRNAs (*P* < 0.05 and |log_2_(fold change)| ≥ 0.5), of which 22 were upregulated and 38 were downregulated in ASD.

### Weighted gene co-expression network analysis (WGCNA)

To build the singed compression network, we used the R package WGCNA [73, 74], a dynamic tree-cutting algorithm, to identify the modules of correlated genes. We followed the similar approach that Wu *et al.* used [15]. In brief, first, pair-wise topology overlap (TO) between circRNAs was calculated with a soft threshold power of 9 to fit scale-free topology. Second, the network was constructed to be robust by performing 1000 rounds of bootstrapping strategy, and the TO matrix was computed for every resampling networks. The median of TOs was used to reconstruct a consensus TO matrix to avoid obtaining the module driven by outliers. Modules were identified using R package dynamicTreeCut with function cutreeDynamic, in which the paramenters are: method = “hybrid”, deepSplit = 3, pamStage = T, pamRespectsDendro = T, minClusterSize = 10, and the expression of each module was summarized by the eigengene (ME), defined as the first principle component of all circRNAs in the module. To identify disease status, the Pearson correlations between MEs and several confounding factors, such as diagnosis, age, sex, brain region, RIN were calculated. Illustration of networks was plotted by the Cytoscape package (https://cytoscape.org/). Finally, to examine the similarity between FC and TC samples, and between ASD and CTL samples, the module preservation analysis was performed using WGCNA package with function modulePreservation. Each module was measured by the Z_summary_ scores, where a module was regarded as “not preserved”, “moderately preserved”, and “highly preserved” if Z_summary_ score < 2, 2 < Z_summary_ score < 10, and Z_summary_ score > 10, respectively. We found that the Z_summary_ scores of all the three DE-modules were greater than or equal to two, indicating that the three modules were preserved across ASD and non-ASD control samples (Fig. 2d) and across FC and TC samples (Fig. 2e).

### Identification of ASD-associated circRNA-miRNA-mRNA regulatory

In this study, we identified 60 DE-circRNAs (22 upregulated and 38 downregulated circRNAs) and three DE-modules (one upregulated module including 21 circRNAs and two downregulated modules including 298 circRNAs) in ASD cortex. To identify the corresponding miRNA-mRNA regulatory axes that were potentially regulated by these ASD-associated circRNAs, we extracted 58 DE-miRNAs (41 upregulated and 17 downregulated miRNAs) in ASD cortex [15] from the study of Wu et al. These ASD-affected miRNAs were derived from 95 human cortex samples (47 samples from 28 ASD cases and 48 samples from 28 non-ASD controls), of which 73 samples overlapped with the samples examined in this study. We then searched for the target sites of the 41 upregulated miRNAs in the 38 downregulated circRNAs and 298 circRNAs defined in two downregulated circRNA modules and the target sites of the 17 downregulated miRNAs in the 22 upregulated circRNAs and 21 circRNAs defined in one upregulated circRNA module, respectively, using RNA22 [75] with default parameters (version 2.0; https://cm.jefferson.edu/rna22/). Next, we identified target genes of the 58 DE-miRNAs. Of the 58 miRNAs, 37 were well-annotated, whereas 21 were newly identified by the study of Wu et al.[15] For the 37 miRNAs, which can be found in the Ingenuity Pathway Analysis (IPA) package [76] (QIAGEN Inc., https://www.qiagenbioinformatics.com/products/ingenuitypathway-analysis) and well-known miRNA databases such as DIANA-TarBase [77], we searched for targets of these miRNAs based on the MicroRNA Target Filter in the IPA package or the latest version of DIANA-TarBase (version 8) [77]. For accuracy, the miRNA-mRNA binding events were considered if they satisfied one of the three criteria: (1) they were TargetScan-precited [78] events with IPA filter score < −0.16 and previously identified by Wu et al. [15]; (2) they were experimentally confirmed events collected by Ingenuity® Expert Findings in IPA, which were manually curated by the IPA experts; and (3) they were experimentally confirmed events collected in DIANA-TarBase (version 8). For the other 21 miRNAs, we identified targets of these miRNAs using the MirTarget algorithm with target prediction scores > 80 (The score was suggested by miRDB, in which the target prediction would be more likely to be real.) via the miRDB [79, 80] Web server at http://mirdb.org/custom.html, where the mature miRNA sequences of the 21 miRNAs were downloaded from the study of Wu et al. [15] For accuracy, we only considered the predicted miRNA-mRNA binding events that were also previously identified by Wu et al. [15] Of note, the predicted miRNA-mRNA binding events previously identified by Wu et al. represented that the binding events should satisfy one of the two criteria (the strongest target and the most conserved target criteria) defined by Wu et al. [15] A total of 10,841 genes were identified to be targets of the 58 DE-miRNAs (see Additional file 5: Table S4). The ASD-associated circRNA-miRNA-mRNA interactions were determined by integrating the circRNA-miRNA interactions with the miRNA-mRNA interactions according to the common miRNA target sites of the circRNAs and mRNAs. We then calculated the correlations between circRNA and miRNA expression, between miRNA and mRNA expression, and between circRNA and mRNA expression based on the same set of cortex samples (i.e., 73 samples). Here, the expression level of the target gene were measured by log_2_ normalized FPKM, which accounted for gene read counts, GC content, gene length, and library size, using the cqn program in the R package [70]. The expression levels of the examined miRNAs were kindly provided by Prof. Daniel H. Geschwind and Ye E. Wu (the authors of the reference [15]), which were measured by log2 normalized read counts for library size, GC content, batch effect, and other technical covariates (RIN, PMI, and batch bank). As shown in Figure 3a, only the circRNA-miRNA-mRNA interactions were considered if they simultaneously satisfied the following rules: (1) both the circRNA-miRNA and miRNA-mRNA interactions should exhibit a significantly negative correlation (one-tailed Spearman’s *P* < 0.05) of expression profile between circRNAs and the corresponding predicted regulated miRNAs and between miRNAs and target mRNAs, respectively; (2) the circRNA expression should be positively correlated with the corresponding mRNA expression; and (3) the Fisher’s combined *P* values [47] of the above three independent Spearman’s correlation tests should be less than 0.05. Finally, 8,170 (including 356 upregulated and 7,814 downregulated circRNA- involved interactions) ASD-associated circRNA-miRNA-mRNA regulatory axes were determined, which included 2,302 target genes. Of note, the experimentally confirmed miRNA-mRNA binding events were obtained according to Ingenuity® Expert Findings in IPA [76] and DIANA-TarBase (version 8) [77].

### Gene set enrichment analysis

The SFARI [3] gene list was downloaded from https://gene.sfari.org/ (Dec. 2018). The high-confidence ASD genetic risk genes (102 genes) were downloaded from the study of Satterstrom et al. [55], which were derived from an enhanced Bayesian analytic framework based on a large dataset of whole-exome sequencing (35,584 ASD subjects). The epilepsy-related gene list was downloaded from the EpilepsyGene database [81] at http://www.wzgenomics.cn/EpilepsyGene/index.php (all epilepsy-related genes). The schizophrenia-related gene lists [82] and the genes associated with human height [83] were downloaded from the GWAS report. The gene lists of Austism_KB genes, iPSD, ePSD, and other brain disorder risk genes were downloaded from the study of Wang, P., et al. [84] at https://www.nature.com/articles/s41398-017-0058-6#Sec16. For Fig. 3c, gene set enrichment analyses were performed using two-tailed Fisher’s exact test with the fisher.test R function. *P* values were False-discovery rate (FDR) adjusted across 14 target groups for each gene list using Bonferroni correction (Fig. 3c). Phenotype ontology analysis was performed using the ToppFunn module of ToppGene Suite software [48]. FDR adjusted *P* values were calculated using Bonferroni correction (Fig. 3b).

### Cell culture

Normal human astrocytes (NHA) cell line was purchased from Gibco (N7805100) and cultured in Human astrocytes growth medium (Cell Applications). Human neuroblastoma (SH-SY5Y) cell line was obtained from ATCC (CRL-2266) and cultured in DMEM/F12 medium (Gibco). Human glioblastoma (U118) cell line was purchased from ATCC (HTB-15) maintained in Dulbecco’s modified Eagle’s medium (Gibco). In addition, human neural progenitor cells (hNPCs; ReNcell VM [85, 86]) were commercially purchased from Sigma-Aldrich (SCC008). The cells were grown on 20μg/ml laminin (Merck) coated culture plates containing ReNCell NSC maintenance medium (Merck) supplemented with 20ng/ml of bFGF and EGF (Merck). All culture media supplemented with 10% fetal bovine serum (FBS) and 1x antibiotic-antimycotic (Gibco) at 37°C with 5% CO_2_. Cells were passaged when the confluence reached 80% of the culture plate every 3 days. Briefly, cells were rinsed with PBS and then incubated in Accutase (Millipore) for 3-5 minutes until cell detached. We used the culture medium to inhibit enzymatic reaction and centrifuged the suspension at 500xg for 5 minutes. We then resuspended the cell pellet in fresh medium and counted the cell number.

### Vector construction and design of oligonucleotide

For overexpress of circARID1A, we constructed the putative exon sequence of circARID1A (E2-E3-E4) into Circle3 vector (provided by Dr. Laising Yen)[87] which can circularized the transcript to produce circular RNA. Putative circARID1A sequence was amplified by PCR, and the PCR product was inserted into the Circle3 vector between Mfe-I and Age-I sites. Sanger sequencing verified recombinant plasmid construction. For ReNcell transfection, cells were infected with lentiviruses which were constructed by National RNAi Core Facility in Academia Sinica (Taipei, Taiwan). The lentiviruses carrying three different vectors: circARID1A-shRNA (pLKO_TRC005), circARID1A-cDNA (pLAS2w.Ppuro), and dual-luciferase with circARID1A cDNA (pLAS2.1w.PeGFP-I2-Puro). All constructs were verified by Sanger sequencing (Additional file 8: Table S7).

### Cell transfection and lentiviral transduction

Cells were transfected with siRNA (MDBio) or miRNA mimics (TOOLS) at final concentration of 10 nM using Lipofectamine RNAiMAX reagent (Invitrogen). The sequences were listed in Additional file 8: Table S7. When transferring plasmids into the cells, the transfection was carried out using TransIT-LT1 Reagent (Mirus) according to the transfection manufacturer’s instructions. For lentiviral transduction, the cells was infected for 24 hours (MOI = 2) and then washed 3 times to remove the remaining virus. After transduction, cells were selected by culture medium containing 0.25μg/ml puromycin for 4 days. The effects of knockdown or overexpression were examined by RT-qPCR. To efficiently knockdown circARID1A expression in different cell types, the siRNA and shRNA targeting the circARID1A backspliced junction were designed. NHA, SH-SY5Y and U118 were transfected with si-circARID1A; and hNPCs were infected with lentiviruses carrying sh-circARID1A.

### Total RNA isolation

Total RNA from cells were extracted using TRIzol reagent with PureLink RNA Mini Kit (Ambion) and PureLink DNase Set (Ambion). Total RNAs of normal human tissues were purchased from Invitrogen. Human brain cDNA array was purchased from Amsbio (HBRT501). Brain total RNA of monkey, mouse and rat were purchased from BioChain. Brain total RNA of rabbit and chicken were purchased from Zyagen. The information of purchased RNA was provided in Additional file 8: Table S7.

### RNase R treatment and subcellular fractionation

For RNase R treatment, total RNA was incubated with or without of RNase R (Epicentre) for 45 minutes at 37 °C to deplete linear and enrich circular RNAs. To validate the subcellular localization preference, we used the NE-PER nuclear and cytoplasmic extraction reagents (Thermo) to separated nuclear and cytoplasmic RNA.

### cDNA synthesis and quantitative Real-Time PCR analysis

For mRNA and circRNA quantitation, RNAs were reverse-transcribed into cDNA using SuperScript III with random hexamer and oligo(dT)_18_ primers (Thermo). qRT-PCR were performed using Luminaris Color HiGreen High ROX qPCR Master Mix (Thermo). Quantity of gene expression is normalized to the expression level of GAPDH. For miRNA quantitation, cDNA synthesis is carried out by or miRCURY LNA RT Kit (Qiagen). qRT-PCR is performed by miRCURY LNA SYBR Green PCR Kit (Qiagen) with miRCURY LNA miRNA PCR assays (Qiagen) for each candidate miRNAs, and U6 small nuclear RNA (snRNA) served as an internal control. All reactions are performed three times per experiment and detected by StepOnePlus Real-Time PCR Systems (Thermo). All primers used in this study were listed in Additional file 8: Table S7.

### Luciferase reporter assays

The luciferase reporter was constructed by subcloning the circARID1A sequence into the Secrete-Pair Dual Luminescence vector (GeneCopoeia). Cells were seeded in 24-well plate and incubated for 24 hours prior to transfection. Dual Luminescence vectors that contained circARID1A sequence were co-transfected into human neuronal cells with miR-204-3p mimics or negative control mimics by TransIT-X2 transfection reagent (Mirus). After 72 hours of transfection, cell culture medium was collected. The luciferase activity was measured by Secrete-Pair Dual Luminescence Assay kit (GeneCopoeia) according to the manufacturer’s protocol. The luciferase activity of Gaussia luciferase (GLuc) was normalized with secreted alkaline phosphatase (SEAP). The fold change was measured by miR-204-3p compared with negative control.

### Neuronal differentiation and Immunostaining

ReNcell VM (cat. no. SCC008, Sigma-Aldrich) was incubated with maintenance medium without containing FGF-2 and EGF growth factors. Maintenance basal media was changed every 3 days for 2 weeks. After 2 weeks, we categorized the neuronal subtypes by immunofluorescence. Cells were plated on glass slides coated with laminin overnight. Cells were fixed with 4% formaldehyde at 37 °C for 25 min, permeabilized with 0.05% Triton X-100 for 15 min, and incubated with 3% FBS for blocking for 1 hr. Cells were then incubated with primary antibody against βIII-tubulin (neuronal marker; cat. no. MA1-118, Invitrogen) or GFAP (glial marker; cat. no. 13-0300, Invitrogen) or nestin (neural stem cell marker; cat. no. MA1-110, Invitrogen) at 4 °C overnight. Next day, cells were incubated with fluorescently labeled secondary antibodies (Additional file 8: Table S7) at 37 °C for 1.5 hours. Nuclei were stained with SlowFade Diamond Antifade Mountant with DAPI (Invitrogen).

### Microarray analysis

The microarray hybridization and the data collection were performed by the Affymetrix GeneChip System Service center in Genomics Research Center, Academia Sinica. Purity and RNA integrity number (RIN) of total RNA was evaluated by 2100 bioanalyzer (Agilent Technologies). Total RNA was hybridized to Affymetrix Human Genome Plus 2.0 Array (Affymetrix). The microarray raw data was analyzed using Transcriptome Analysis Console (TAC 4.0) software (Affymetrix).

### Availability of data and materials

The code for the DE-circRNA analysis and the related input data were publicly accessible at GitHub (https://github.com/TreesLab/lme_DEcircRNA). Numbers of the detected circRNAs per million mapped RNA-seq reads in the ASD and non-ASD control samples from the three brain regions (FC, TC, and CV) are provided in Additional file 1: Table S1. The 53,427 identified circRNAs in the 202 cortex samples (73 FC, 61 TC, and 68 CV samples), including the coordinates of the circRNA junctions and the corresponding circRNA expression levels (*RPM*), are deposited in Additional file 2: Table S2. The 1,060 identified circRNAs detected in more than 50% of the 134 cortex samples are deposited in Additional file 3: Table S3. The 10,841 identified targets of the 58 DE-miRNAs are provided in Additional file 5: Table S4. The 8,170 identified ASD-associated circRNA-miRNA-mRNA regulatory axes and the 2,302 target genes of the circRNA-miRNA-mRNA axes are provided in Additional file 6: Table S5. Microarray results of the fold changes for the 158 target genes of the circARID1A-miR-204-3p-involved interactions with the treatments of circARID1A and miR-204-3p overexpression are provided in Additional file 7: Table S6. All primers and antibodies used in this study are provided in Additional file 8: Table S7.

## Additional files

**Additional file 1: Table S1.** Numbers of the detected circRNAs per million mapped RNA-seq reads in the ASD and non-ASD control samples from the three brain regions (FC, TC, and CV). (XLSX 17 kb)

**Additional file 2: Table S2.** The 53,427 identified circRNAs in the 202 cortex samples (73 FC, 61 TC, and 68 CV samples). The coordinates of the circRNA junctions and the corresponding circRNA expression levels (*RPM*) are deposited in the online file. (XLSX 38193 kb)

**Additional file 3: Table S3.** The 1,060 identified circRNAs that were detected in more than 50% of the 134 cortex samples examined. The coordinates of the circRNA junctions and the related information are deposited in the online file. (XLSX 1130 kb)

**Additional file 4: Figure S1.** Resampling analysis with 100 rounds of random sampling of 70% of the samples examined. **Figure S2.**Numbers of the predicted binding sites of the previously identified upregulated (or downregulated) miRNAs in ASD on the circle sequences of the identified downregulated (or upregulated) circRNAs/circRNA modules. **Figure S3.** The 7,814 upregulated circRNA-involved circRNA-miRNA-mRNA interactions plotted by the Cytoscape package. **Figure S4.** RT-PCR experiments representing that circARID1A is widely expressed in 24 human brain regions. **Figure S5.** The circRNA-mRNA and miRNA-mRNA correlations of expression profile between circARID1A and the 171 target mRNAs (red line) and between miR-204-3p and the 171 target mRNAs (green line), respectively. **Figure S6.** Enrichment of high-confidence ASD risk genes for the targets of the identified circRNA-miRNA-mRNA interactions. **Figure S7.**Principal component analysis (PCA) based on the overall circRNA expression profiles of all identified circRNAs (1,060 circRNAs) from the cortex samples. (DOCX 1832 kb)

**Additional file 5: Table S4.** The 10,841 identified targets of the 58 DE-miRNAs. (XLSX 1364 kb)

**Additional file 6: Table S5.** The 8,170 identified ASD-associated circRNA-miRNA-mRNA regulatory axes and the corresponding target genes of the circRNA-miRNA-mRNA axes. The related information was also provided. (XLSX 1605 kb)

**Additional file 7: Table S6.** Microarray analysis of the fold changes for the 171 target genes of the circARID1A-miR-204-3p-involved interactions with the treatments of circARID1A and miR-204-3p overexpression. (XLSX 16 kb)

**Additional file 8: Table 7.** The primers and antibodies used in this study. (XLSX 13 kb)

## Acknowledgements

We thank the National RNAi Core Facility (Academia Sinica) for providing shRNA reagents and related services and Affymetrix Facility for microarray analysis (Genomics Research Center (GRC), Academia Sinica), respectively. We especially thank Profs. Michael Hsiao, Ying-Chih Chang, and Jean Lu for providing experimental assistance and Prof. Daniel H. Geschwind and Ye E. Wu for providing the related data of miRNAs examined.

## Author contributions

CYC provided code for the analyses of DE-circRNAs and circRNA-miRNA-mRNA regulatory interactions. TLM performed the analysis of circRNA coexpression networks. YJC and CYC collected data. YJC performed DE-circRNA identification and most experimental validations of the study. CFC and YDW gave assistance to experiments. SKG and LY provided the plasmid for circRNA overexpression. TJC conceived the project, designed the full research framework, analyzed data, and wrote the manuscript.

## Ethics approval and consent to participate

The rRNA-depleted RNA sequencing data of human brain was obtained by the request from Synapse (http://www.synapse.org) under the accession number syn4587609. All the cell lines and total RNAs of normal human/animal tissues used in this study are commercialized samples. Ethics approval for the use of these samples was granted to TJC by the Ethics Committee of Academia Sinica (no. BSF0418-00003896).

